## Supplementary Figures for "Longitudinal Long-Read Microbiome Profiling in a Canine Model Reveals How Age, Diet, and Birth Mode Shape Gut Community Dynamics"

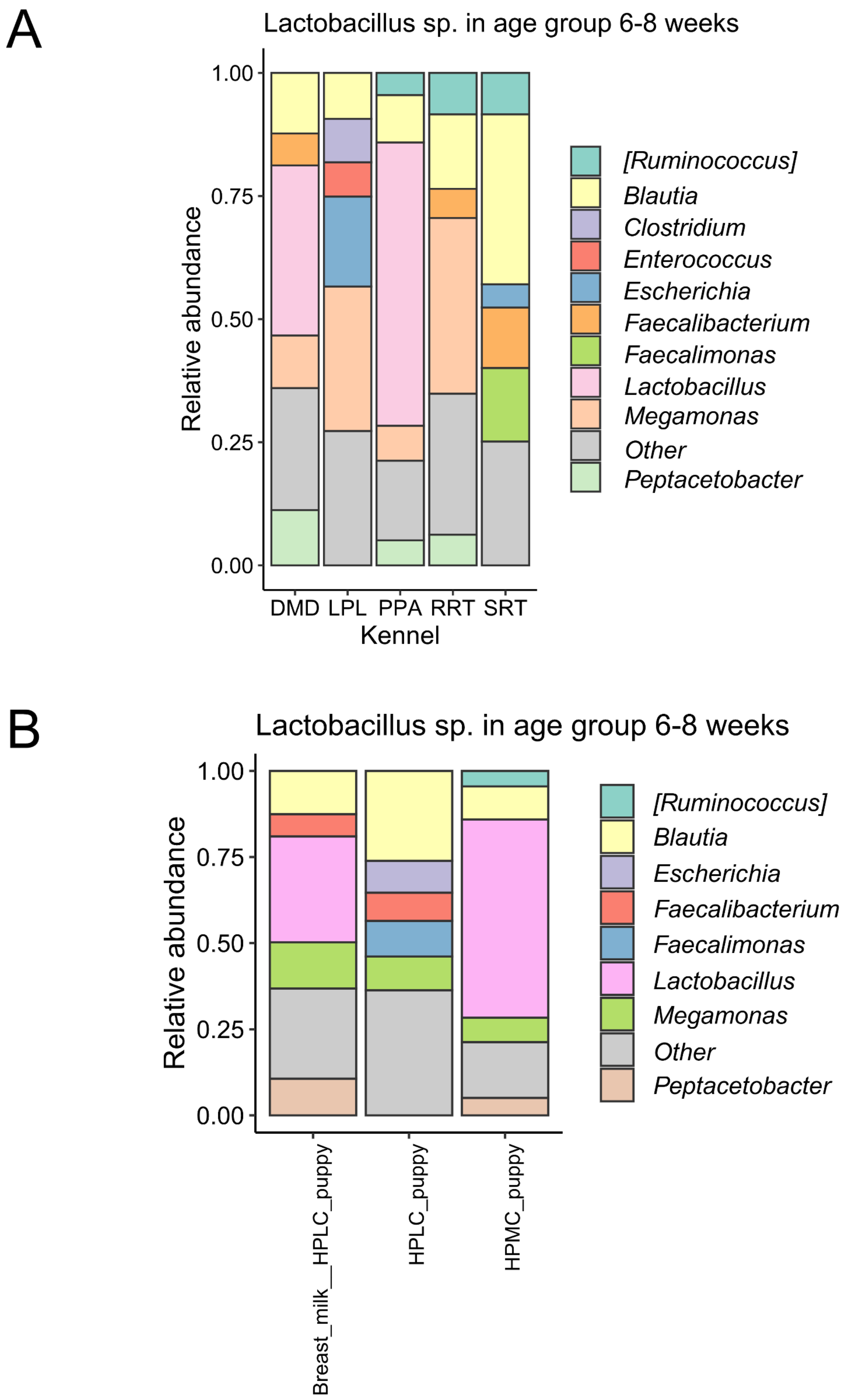


**Fig. S1. Early-Weaning Lactobacillus Patterns (6–8 Weeks): Kennel and Diet.**
**A)** Relative abundance of the top 10% most abundant genera, focusing on *Lactobacillus sp.* Presence in the 6-8 weeks age group per kennel. **B)** *Lactobacillus sp.* Abundance per diet in the 6-8 weeks age group.

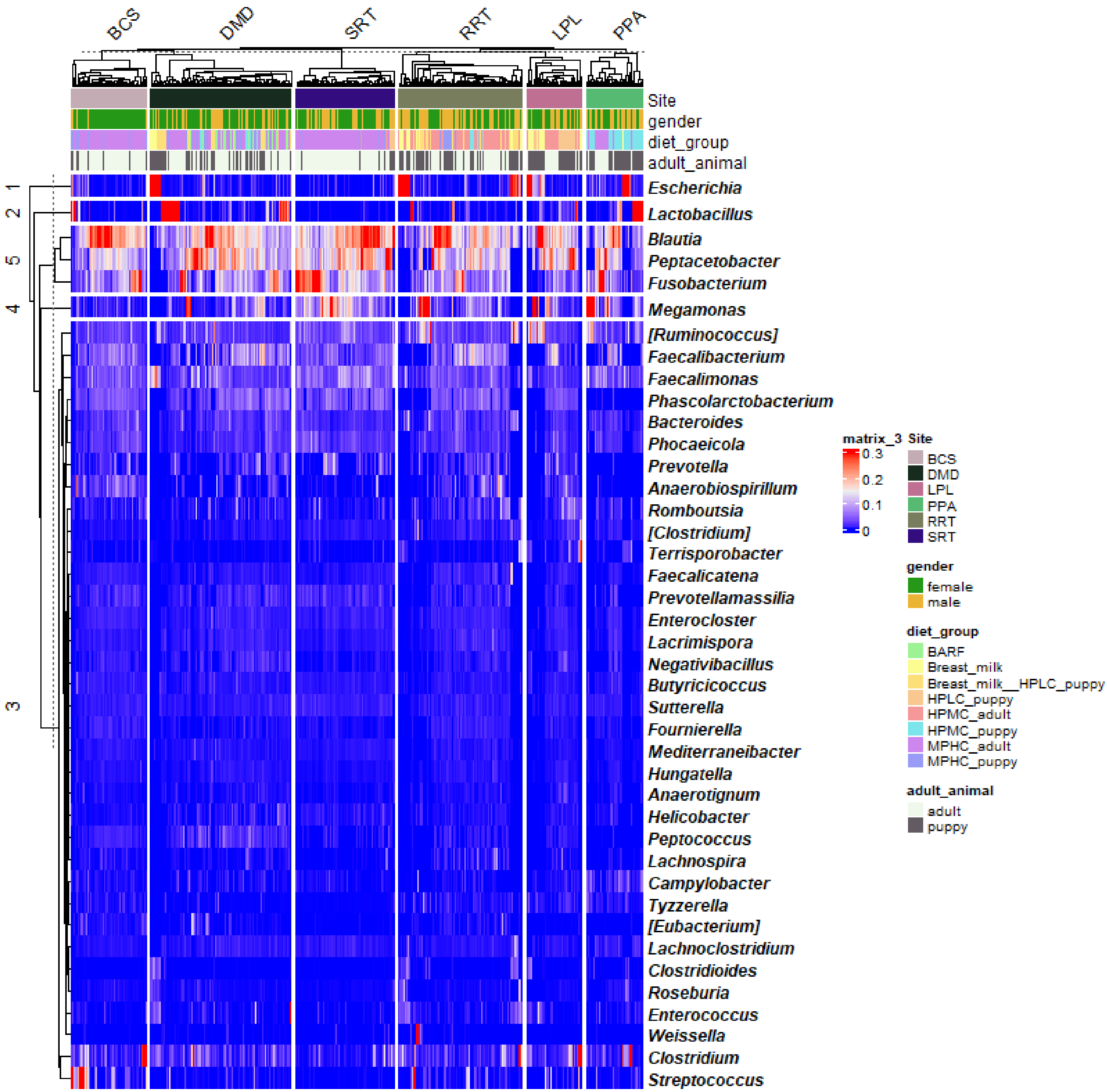


**Fig. S2. Heatmap of relative abundances of the top 40 genera across all kennels.**
Hierarchical clustering shows that age and diet provide more structured clustering than sex. While *Blautia* is represented by several different species in the dataset, complicating the analysis, genus-level clustering results in characteristic patterns for this taxon.

A

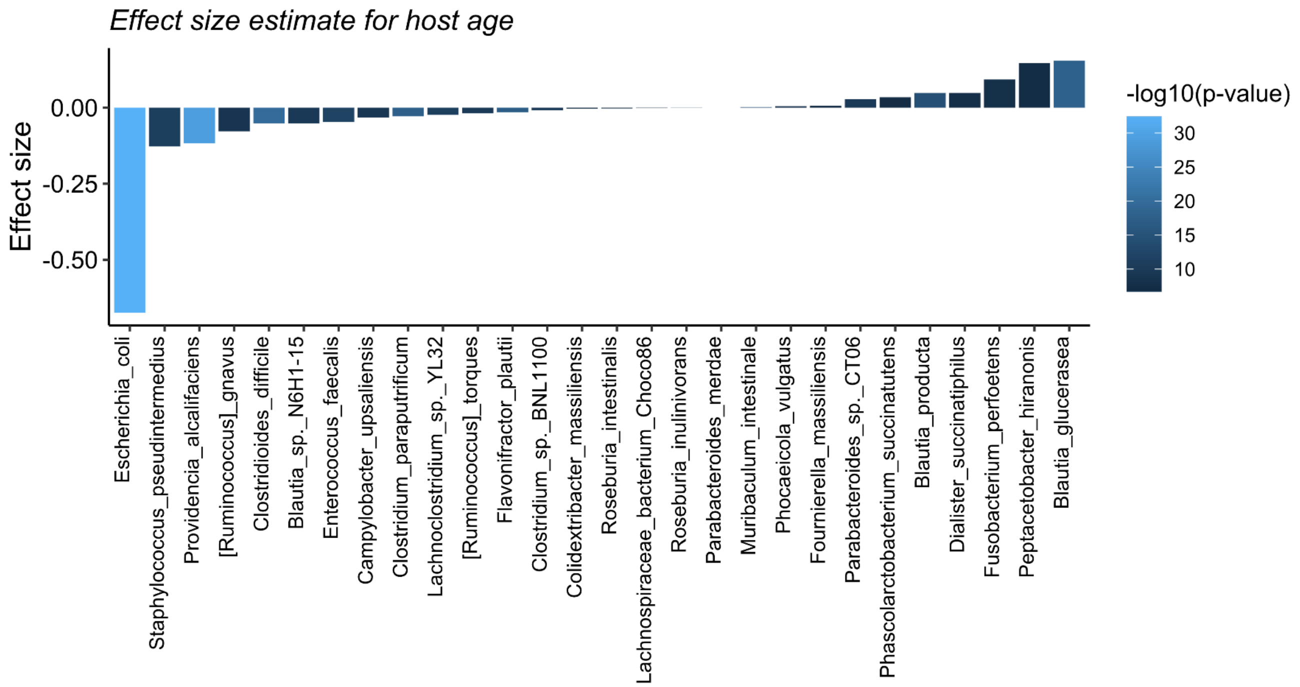


B

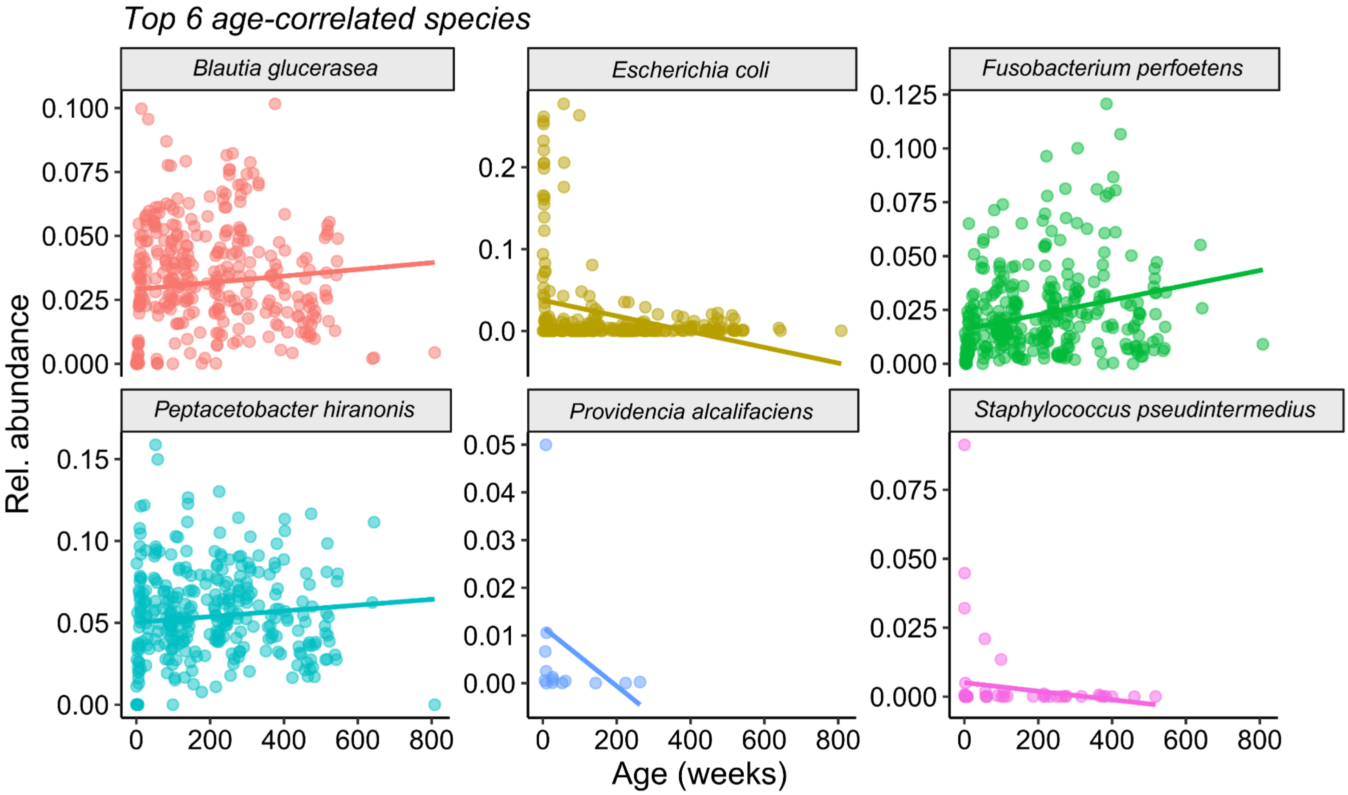


**Fig. S3. Age-Associated Species: Effect Sizes and Abundance Trajectories.**
**A)** Bar chart of mixed-effects model estimates for age-correlated species. Y-axis displays the effect size estimate, color gradient is proportional to p-value of the interaction. **B)** Scatter plots of species relative abundance vs. host age in weeks; best-fit lines indicated per species.

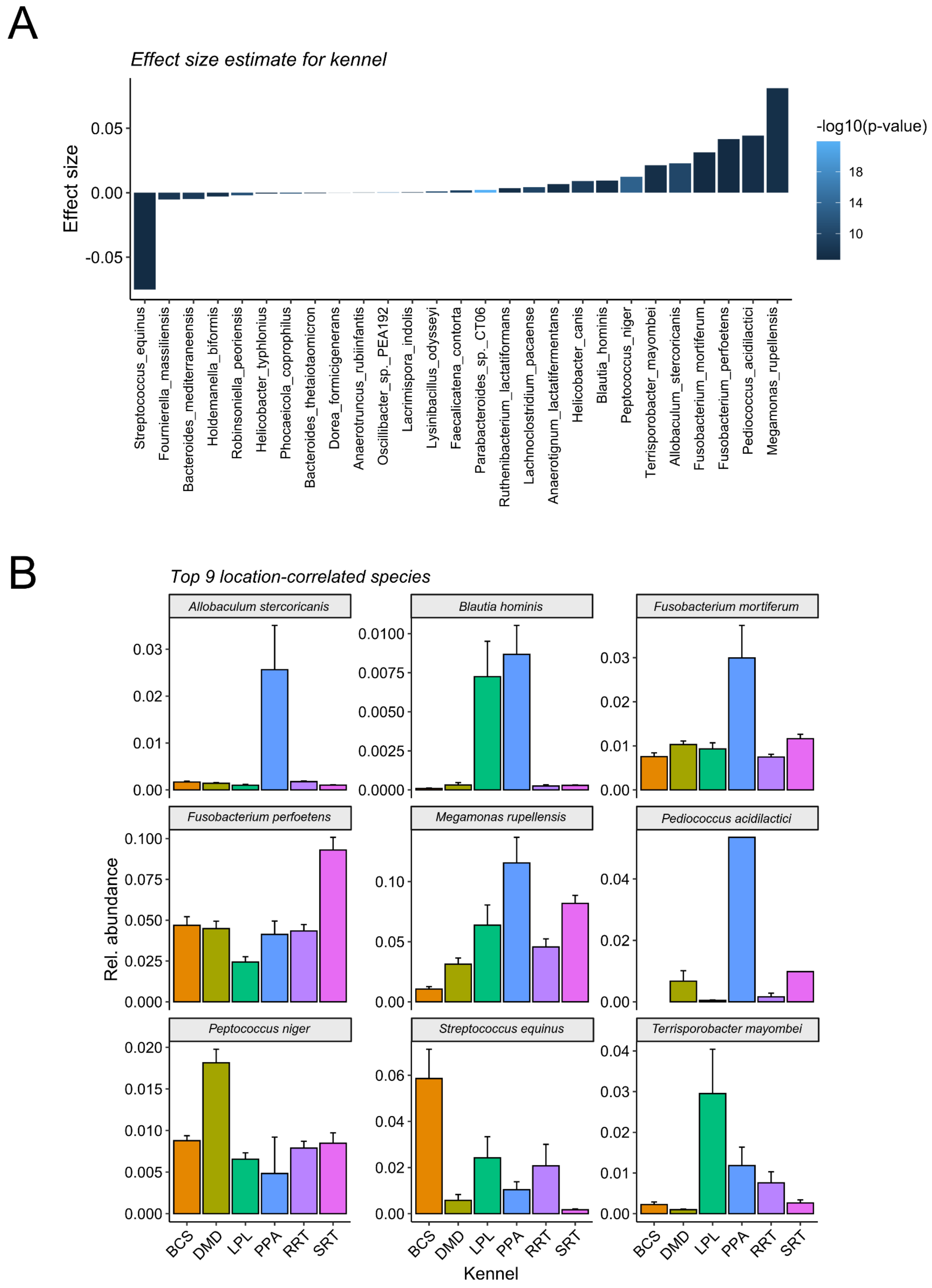


**Fig. S4. Kennel-associated microbial composition.**
**A)** Bar chart of mixed-effects model estimates for kennel-correlated species. Y-axis displays the effect size estimate, color gradient is proportional to p-value of the interaction. **B)** Bar charts of species relative abundance vs. location; error bars indicate standard deviation.

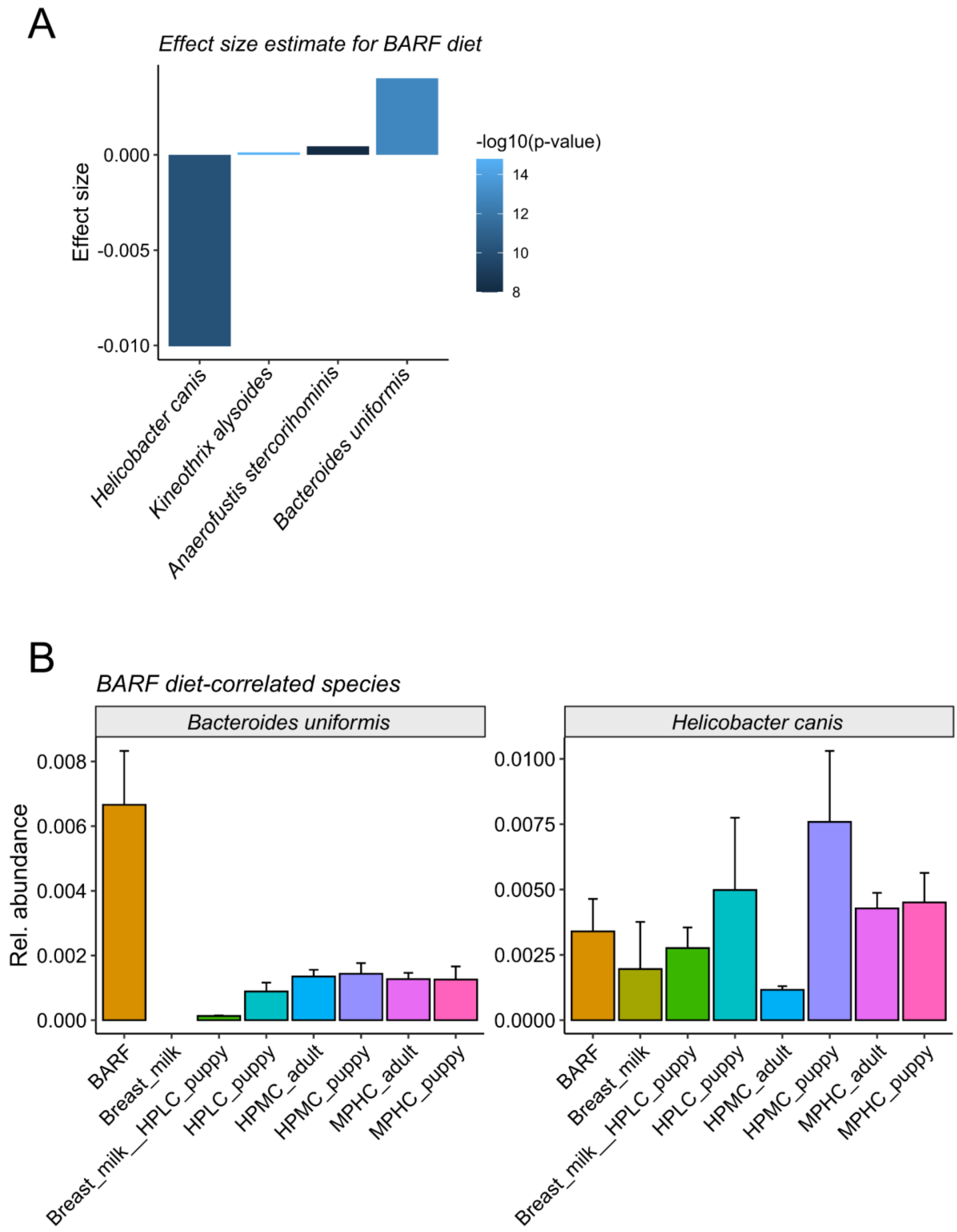


**Fig. S5. Effects of BARF-diet on Microbial Composition.**
**A)** Effect size estimates from mixed-effects models for diet-associated species.Y-axis displays the effect size estimate, color gradient is proportional to p-value of the interaction. **B)** Bar charts of species relative abundance vs. Diet category; error bars indicate standard deviation. Abbreviations for host diet terms: HPLC (high-protein–low-carbohydrate), HPMC (high-protein–moderate-carbohydrate), MPHC (moderate-protein–high-carbohydrate).

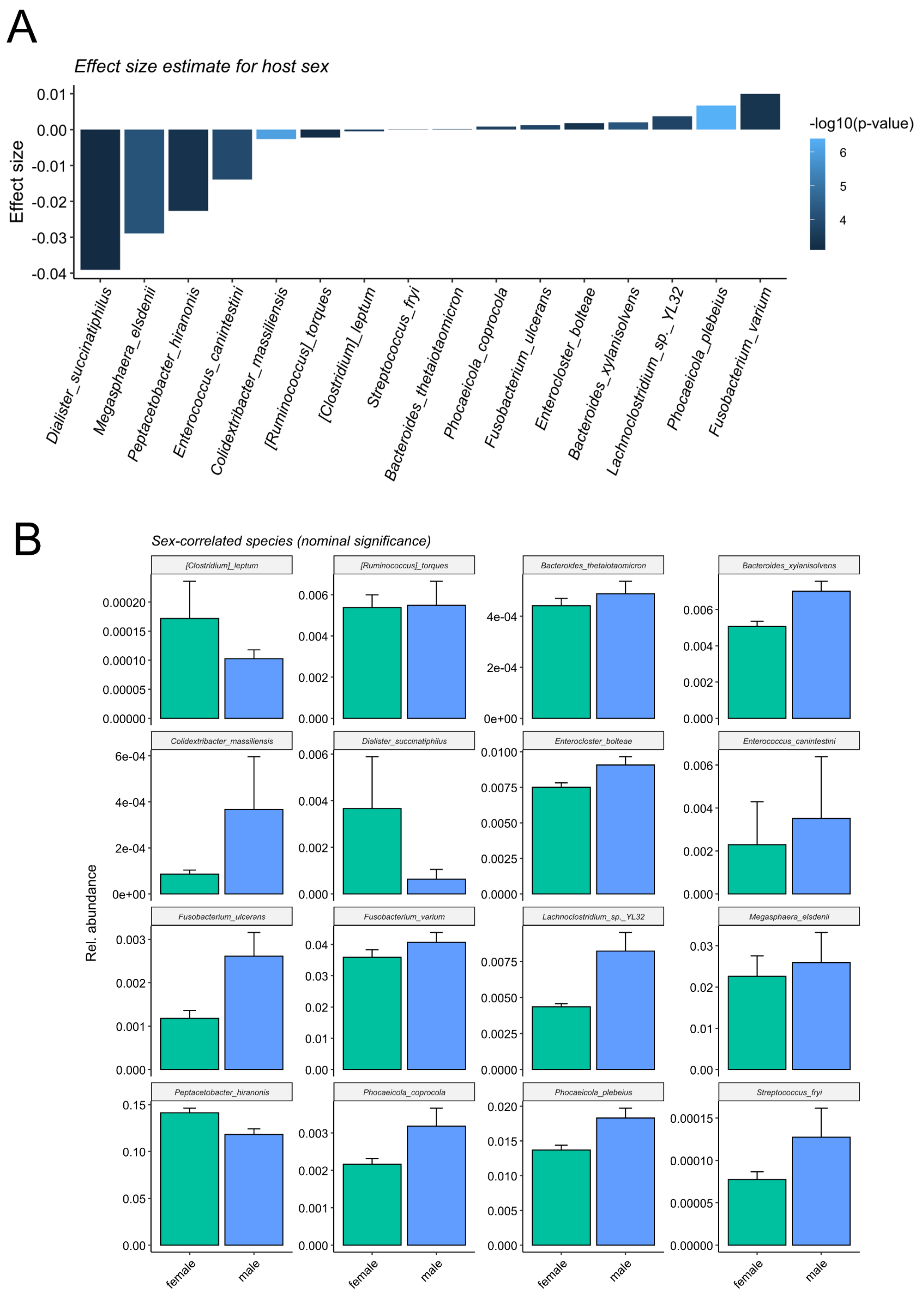


**Fig. S6. Sex-Associated Microbial Composition.**
**A)** Bar chart of mixed-effects model estimates for sex-associated species. Y-axis displays the effect size estimate, color gradient is proportional to p-value of the interaction. **B)** Bar charts of species relative abundance vs. sex; error bars indicate standard deviation.
