## Supplementary File for "Longitudinal Long-Read Microbiome Profiling in a Canine Model Reveals How Age, Diet, and Birth Mode Shape Gut Community Dynamics"

### SUPPLEMENTARY METHODS

#### DNA extraction – detailed protocol

Fecal samples stored at −80 °C were thawed at room temperature for 40 min. Each sample was homogenized prior to aliquoting to ensure even microbial composition and to minimize microgeographical variation. Genomic DNA was extracted using the ZymoBIOMICS™ 96 MagBead DNA Kit-Dx (Zymo Research, Cat. No. D4308-E) according to the manufacturer's instructions, under sterile conditions in a Class II biosafety cabinet (SafeMate EZ 1.2, BIOAIR). For each sample, 750 µl of stool was transferred into a ZR BashingBead™ Lysis Tube (0.1 & 0.5 mm) and subjected to bead beating on a Vortex-Genie® 2 platform (Scientific Industries) for 40 min at speed 10. The lysate was centrifuged at 10,000 × g for 1 min, and 200 µl of the supernatant was transferred into a 1.5 ml DNA LoBind microtube (Eppendorf™ Safe-Lock). Next, 600 µl of ZymoBIOMICS™ MagBinding Buffer and 25 µl of ZymoBIOMICS™ MagBinding Beads were added. The mixture was pipette-mixed five times and incubated on a shaker plate (Vortex-Genie® 2, Scientific Industries) at speed ≤2 for 10 min. Tubes were placed on a magnetic stand (MagJET Separation Rack, Thermo Fisher Scientific) until beads pelleted. The supernatant was removed, and the pellet was washed with 500 µl of MagBinding Buffer, then with 500 µl of ZymoBIOMICS™ MagWash 1, and twice with 900 µl of ZymoBIOMICS™ MagWash 2, with each wash involving 1 min of mixing followed by brief centrifugation. Beads were air-dried by incubation at 55 °C for 10 min on a dry block heating element (Bio TDB-100, bioSan). DNA was eluted in 50 µl of ZymoBIOMICS™ DNase/RNase-Free Water by pipette mixing (5×) and incubating for 10 min on a shaker plate, followed by bead separation on a magnetic stand for 2–3 min. The eluate was transferred into a new DNA LoBind tube. DNA concentration was measured with a Qubit® 4 Fluorometer using the Qubit dsDNA HS Assay Kit (Invitrogen, USA), and DNA integrity was assessed on an Agilent 4150 TapeStation with Genomic DNA ScreenTape. gDNA was stored at −20 °C until library preparation.

**16S library preparation – detailed protocol**

High-quality genomic DNA (DIN ≥ 6) was diluted to 7.5 µl with nuclease-free water (ThermoFisher Scientific, USA) and combined with 12.5 µl of LongAmp® Hot Start Taq 2X Master Mix (New England Biolabs, USA) and 5 µl of 16S barcode from the Oxford Nanopore Technologies (ONT) Rapid Sequencing Amplicons–16S Barcoding Kit (SQK-16S024). The mixture was gently pipette-mixed on ice.

PCR amplification was performed on a ProFlex PCR System (Applied Biosystems, Life Technologies) with the following cycling conditions:

- Initial denaturation at 95 °C for 1 min
- 25 cycles of: 95 °C for 20 s, 55 °C for 30 s, 65 °C for 2 min
- Final extension at 65 °C for 5 min, hold at 4 °C.

At room temperature, 15 µl of AMPure XP beads (Beckman Coulter, USA) was added to the PCR product and incubated for 5 min on a shaking mixer platform (Multi Bio RS-24, bioSan). Tubes were placed on a magnetic rack, the supernatant was discarded, and beads were washed twice with 100 µl of freshly prepared 70% ethanol. Beads were air-dried for 30 s and resuspended in 11 µl of Tris-HCl buffer (10 mM Tris-HCl, 50 mM NaCl, pH 8.0; bioWORLD, USA), incubated for 2 min, and placed back on the magnetic rack to recover the eluate.

Library concentration was determined using a Qubit® 4 Fluorometer with the Qubit dsDNA HS Assay Kit. Fragment size and quality were assessed on an Agilent 4150 TapeStation with D5000 ScreenTape, confirming the expected amplicon size (~1500 bp, full-length 16S V1–V9).

**Sequencing details**

Twenty-four metabarcoded libraries were pooled equimolarly to a final amount of 100 fmol and sequenced on an ONT MinION platform using R9 flow cells. Sequencing data were basecalled with Dorado v0.8.0 in super accurate mode (minimum Q-score = 10).
